# Alignment-free Comparison of Metagenomics Sequences via Approximate String Matching

**DOI:** 10.1101/2020.05.24.113852

**Authors:** Jian Chen, Le Yang, Lu Li, Steve Goodison, Yijun Sun

## Abstract

Quantifying pairwise sequence similarities is a key step in metagenomics studies. Alignment-free methods provide a computationally efficient alternative to alignment-based methods for large-scale sequence analysis. Several neural network-based methods have recently been developed for this purpose. However, existing methods do not perform well on sequences of varying lengths and are sensitive to the presence of insertions and deletions. In this paper, we describe the development of a new method, referred to as AsMac, that addresses the aforementioned issues. We proposed a novel neural network structure for approximate string matching for the extraction of pertinent information from biological sequences and developed an efficient gradient computation algorithm for training the constructed neural network. We performed a large-scale benchmark study using real-world data that demonstrated the effectiveness and potential utility of the proposed method. The open-source software for the proposed method and trained neural-network models for some commonly used metagenomics marker genes were developed and are freely available at www.acsu.buffalo.edu/~yijunsun/lab/AsMac.html.

## 1 Introduction

Microbes play an essential role in processes as diverse as those related to human health and biogeochemical activities critical to life in all environments on earth. However, due to the inability of traditional techniques to cultivate most microbes, our understanding of complex microbial communities is limited. The advent of next-generation sequencing technology allows researchers to study genetic materials recovered directly from natural environments and thereby opens a new window to extensively probe the hidden microbial world [1]. Consequently, metagenomics, where the amplicon sequencing of prokaryotic marker genes (e.g., 16S and 23S rRNA genes) serves as a major probing tool, has recently become an exploding research area [2] and was selected as one of the ten technical breakthroughs in 2013 by the Science magazine [3].

A key step in metagenomics analyses is to quantify pairwise sequence similarities, which plays a fundamental role in various bioinformatics methods for database search, sequence annotation, and sequence binning [4--8]. Alignment distances are generally considered the gold standard; however, due to the high computational complexity, alignment-based methods can only be applied to small sequence datasets. In a metagenomics study, tens of millions or even hundreds of millions of sequences are typically generated. In this case, alignment-free methods [9--12] are perhaps the only computationally viable approach to estimating pairwise sequence similarities. Commonly used alignment-free methods can be broadly classified into two categories: 1) methods based on word frequency (e.g., *k*-mer [13], FFP [14], and CV [15]), and 2) methods based on substrings (e.g., ACS [16], Kr [17], and kmacs [18]). There also exist some methods based on information theory (e.g., IC-PIC [19]), but they are less commonly used. By transforming sequences into numerical vectors, alignment-free methods can be viewed as defining an implicit mapping function that projects sequences into an embedding space where pairwise sequence similarities are calculated. The methods in the first category are based on the statistics of fixed-length word frequency or on the information content of word frequency distributions, while the methods in the second category employ the similarities or differences of substrings in a pair of sequences. For substring-based methods, it is not necessary to specify a word length, and thus in general they can achieve better performance than those relying on a fixed word length. However, substring-based methods introduce new parameters (e.g., the number of mismatches in kmacs [18]) that cannot be easily estimated. Moreover, computing features of variable word lengths usually requires more complex data structures and thus is computationally more expensive [18]. A major limitation of the above methods is that they are all data-independent approaches, where distance measures are predefined based on various heuristics and thus can only provide rough approximations to alignment distances.

Recently, several attempts have been made to use neural networks to develop data-dependent approaches for alignment-free sequence comparison (e.g., SENSE [20] and NeuroSEED [21]). The basic idea is to use a neural network to learn an explicit mapping function through training and map sequences onto an embedding space so that the mean square error between alignment distances and pairwise distances defined in the embedding space is minimized. Since the mapping function is learned through training, it has been demonstrated that data-dependent methods can offer much more accurate solutions than data-independent counterparts [20, 21]. Moreover, neural network-based approaches are computationally very efficient, and run even faster than the *k*-mer method [20]. However, existing methods suffer from several major limitations. First, biological sequences under comparison usually have varying lengths, however, SENSE can only be applied to sequences of equal length, and NeuroSEED uses zero padding to make sequences have the same length, which is not biologically meaningful and can lead to poor performance. Second, early attempts used the convolutional neural network (CNN) for sequence comparison. CNN was designed mainly for image analysis [22] and cannot effectively model insertions and deletions—collectively called indels—due to the use of dot product to measure the similarity between a filter and a sequence. Although gated recurrent units and attention transformers have achieved great success for natural language data analysis [23,24], it was found that they were inferior to CNN for biological sequence comparisons [21].

**Figure 1:**
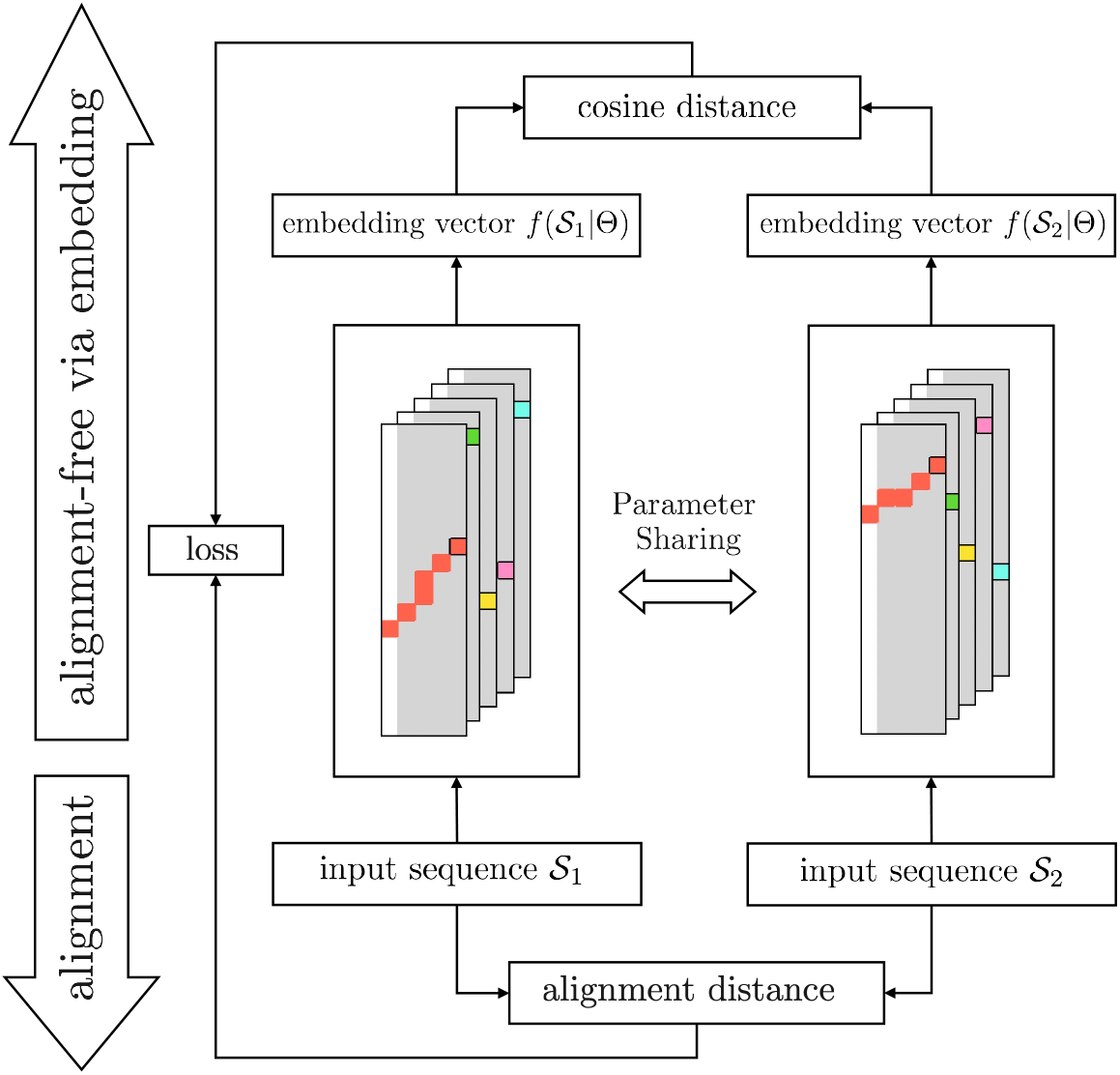
Overview of the proposed method that combines a Siamese neural network and approximate string matching for alignment-free sequence comparison and its training process.

In this paper, we describe the development of a novel neural network-based method, referred to as AsMac, that addresses the aforementioned issues. Specifically, we used approximate string matching (ASM) to transform sequences into numeric vectors. Since ASM uses the edit distance to measure the similarity between a pattern and a sub-sequence, it is well suited for extracting pertinent information from biological sequences. To learn patterns from data *auto-matically*, we proposed a novel neural-network structure for the implementation of approximate string matching and an efficient gradient computation algorithm for training the constructed neural network. We performed a large-scale experiment on prokaryotic rRNA sequence datasets that demonstrated the effectiveness and utility of the proposed method. We also provided trained neural-network models for some commonly used prokaryotic marker genes in an accompanying website that researchers can use directly to process their own datasets.

## 2 Methods

### 2.1 Overview

We propose a novel method that uses a Siamese neural network and approximate string matching for alignment-free sequence comparison. Fig. 1 depicts the overview of the proposed method. The basic idea is to use a neural network to project sequences into embedding vectors so that the difference between alignment distances and pairwise dissimilarities calculated in the embedding space is minimized. Since inputs and outputs are from the sequence and numerical domains, respectively, and are not directly comparable, we use a twin neural network—also called Siamese neural network [25]—to learn a mapping function. Specifically, given a pair of sequences 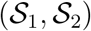, each network takes one sequence as input and outputs 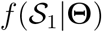 and 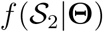 as the embedding vectors of the two sequences, respectively. Here, *f* is the mapping function, and **Θ** is the parameters of the network to be optimized. To form the twin network, we propose a novel neural network for approximate string matching (detailed below). To train the network, we compute the alignment distance 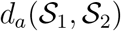 by using the Needleman-Wunsch (NW) algorithm and the cosine distance of the embedding vectors as the embedding distance 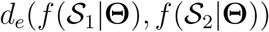, and optimize the neural network by using backpropagation to minimize the mean square error given by

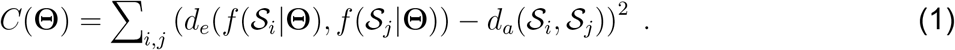

In the training process, the twin networks are forced to share the same parameters, including both initialization and gradient descent updates. Once Siamese neural network is optimized, one of the networks is used to project sequences into embedding vectors.

### 2.2 Approximate String Matching

We start by giving a brief description of the approximate string matching (ASM) algorithm. Let 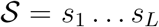 be a sequence and 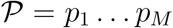 be a pattern. We aim to identify a sub-sequence in 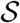 that has the minimum edit distance to pattern 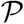 among all subsequences of 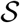. Dynamic programming provides the optimal solution for this purpose [26]. Specifically, we first construct a scoring matrix **F** of size (*M* + 1) × (*L* + 1), where the (*m*, ℓ)-th entry *F_m,ℓ_* is the negative minimum edit distance between the first *m* characters of 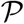 and any sub-sequence 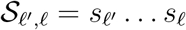 of 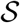 that ends at position ℓ. The scoring matrix can be constructed recursively:

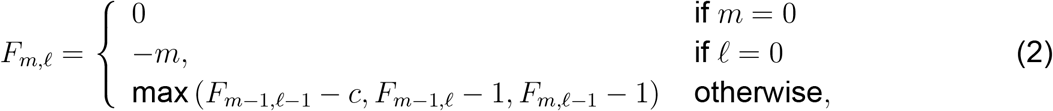

where *c* is the substitution cost that takes a value of 0 if *p_m_* = *s_ℓ_* and 1 otherwise. Once **F** is computed, the maximum value in the last row is the optimal score, and the best-matched subsequence can be identified through backtracking. The computational complexity is on the order of 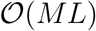. Since pattern 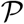 is usually much shorter than sequence 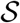, the scoring matrix can be computed efficiently. Moreover, unlike CNN, approximate string matching uses the edit distance to measure the similarity between a pattern and a sub-sequence, *allowing for deletions and insertions*. Thus, it is well suited for extracting pertinent information from biological sequences.

### 2.3 Novel Neural Network for Approximate String Matching

A major issue with approximate string matching for our application is that patterns of interest are generally unknown and can only be pre-determined heuristically. To address the issue, we propose a novel neural network for approximate string matching that enables the learning of patterns from data *automatically*. To cast the problem into a continuous optimization problem so that the constructed neural network can be trained through gradient descent, some modifications are merited. First, we use one-hot coding [20] to transform an input sequence S into an *L* × 4 matrix 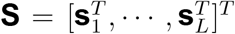, and pattern 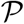 into an *M* × 4 matrix 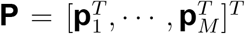. Here, each element in **P** can be learned through training (detailed below). Second, we replace the substitution cost *c* with 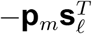 to measure the cost of the mismatch between the m-th vector of the pattern and the ℓ-th vector of the input sequence. Third, while in the original algorithm the gap penalty is set to 1, in order to make the constructed neural network more generally applicable, following the work of [27], we set it to be a learnable parameter. Finally, in order for the constructed mapping function to be differentiable, we replace the max function in Eq. (2) with a generalized maximization operator [28], given by

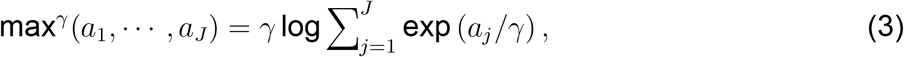

where *γ* > 0 is a smoothing parameter. With the above modifications, Eq. (2) can be reformulated as:

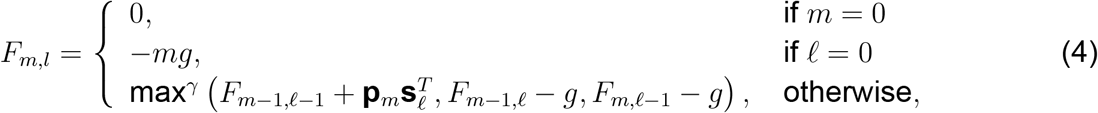

where *g* is the gap penalty. Using the above-modified approximate string matching, we construct a neural network to map input sequences into embedding vectors. Specifically, given an input sequence 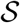 and *N* patterns 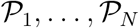, we generate *N* scoring matrices **F**_1_,…,**F**_*N*_. We extract the maximum value *v_n_* from the last row of each scoring matrix **F**_*n*_, and concatenate them into vector **v** to compute an embedding vector

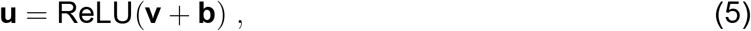

where ReLU is the rectifier activation function [29] and **b** is a bias term.

### 2.4 Learning Parameters through Backpropagation

Finally, we discuss how to estimate the parameters of the proposed method, specifically the patterns used in approximate string matching and the gap penalty, efficiently through backpropagation by minimizing the mean squared error calculated by comparing embedding distances and alignment distances (see Eq. (1)). While backpropagation can be easily implemented by using PyTorch [30], the computation of the gradient of the mean squared error with respect to the patterns is not trivial. Since the patterns are independent of each other, we only need to consider one pattern. Specifically, given pattern matrix **P**, we need to calculate the partial derivative of the mean squared error with respect to each row vector of the pattern matrix

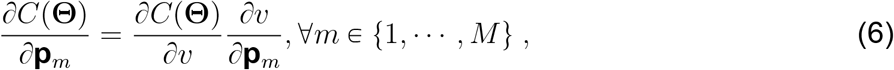

where *v* is the maximum value of the last row of scoring matrix **F**. We note that while the computation of **F** depends on an input sequence, only the best-matched sub-sequence of the input sequence is involved in the computation of *v*. We utilize this fact to develop an efficient scheme for the computation of gradients. By definition, *v* is the score of the global alignment between the pattern and its best-matched sub-sequence, and thus can be computed by using the Needleman-Wunsch (NW) algorithm [31]. An issue with the NW method is that its output is not differentiable. To address the issue, we resort to the differentiable Needleman-Wunsch algorithm [27] to approximate the value of *v* and compute the partial derivatives with respect to the pattern and the gap penalty. Let 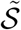 be the best-matched sub-sequence, and 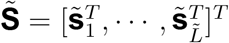. We construct an 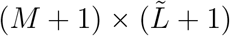 scoring matrix **F** for the differentiable Needleman-Wunsch algorithm:

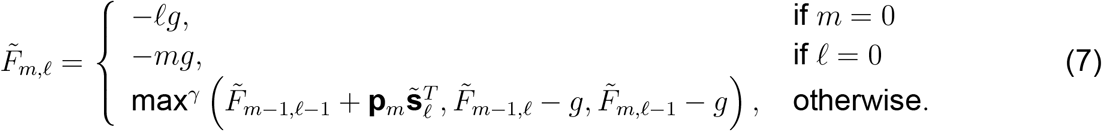

By construction, 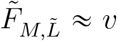. Thus, the approximation of the partial derivative of *v* with respect to each row of pattern **P** can be computed as:

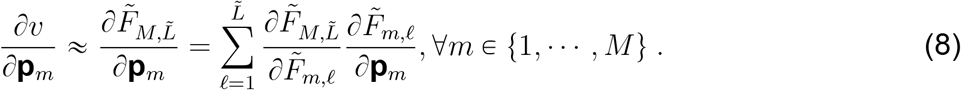

It can be shown that 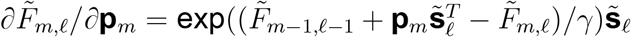. Note that 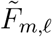 is involved in the calculation of its three neighbors 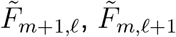 and 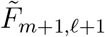. Hence, 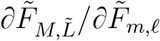 can be computed recursively. Specifically, when *m* ≠ *M* and 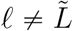,

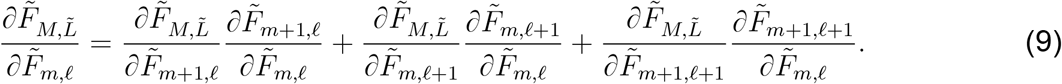

Likewise, the partial derivative *∂v*/*∂g* can be approximated as 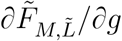, which can be calculated recursively by taking the partial derivative of Eq. (7) with respect to *g*. Specifically, when *m* ≠ 0 and ℓ ≠ 0,

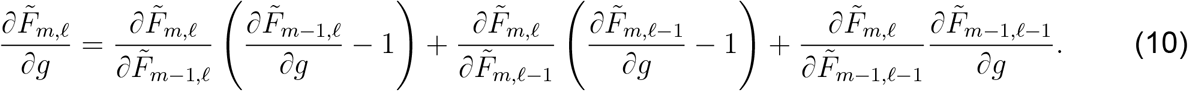

## 3 Experiments

We performed a large-scale benchmark study to demonstrate the effectiveness and potential utility of the proposed method.

**Table 1:** Summary of the six sequence datasets used in the study.

| Dataset | # samples | # reads | sequence length | sequence type |
| --- | --- | --- | --- | --- |
| Qiita | 66 | 6,734,572 | 151 | 16S V4 |
| RT988 | 90 | 4,119,942 | 464-465 | 16S V3-V4 |
| Greengenes | N/A | 925,718 | 1,325-1,500 | 16S full |
| Zymo | N/A | 69,367 | 1,187-1,518 | 16S full |
| Labonte lake | 16 | 41,647 | 385-410 | 23S V5 |
| Silva-23S | N/A | 39,176 | 2,750-3,150 | 23S full |

### 3.1 Sequence Datasets

Table 1 describes the six sequence datasets used in the study. The Qiita dataset was generated from 66 skin, saliva, and fecal samples collected from the Amerindian Yanomami people [32]. It contains 6,734,572 sequences of 151 bps in length, covering the V4 hyper-variable region of the prokaryotic 16S rRNA gene. The RT988 dataset contains 4,119,942 sequences from 90 oral plaque microbiome samples [33]. The sequences have a length of 464-465 bps, covering the V3-V4 hyper-variable region of the 16S rRNA gene. Both datasets were generated by Illumina MiSeq, and before analysis, pre-processing comprised of pair-end joining, quality filtering, and length filtering was performed. The Labonte lake dataset was generated from water samples collected from a eutrophic lake [34], which contains 41,647 sequences with a length ranging from 385 to 410 bps and covering the V5 region of the 23S rRNA gene. The Greengenes dataset was extracted from the Greengenes database [35] and contains 925,718 unique full-length 16S rRNA gene sequences with a length ranging from 1,325 to 1,500 bps. The Silva-23S dataset was downloaded from the Silva database [36], containing 39,176 full-length 23S rRNA gene sequences with a length ranging from 2,750 to 3,150 bps. The Zymo dataset was generated from a Zymo mock community sequenced by PacBio circular consensus sequencing technology [37]. We used the *removePrimers* function from the DADA2 R package [38] to remove primers and orient all the sequences in the forward direction. After pre-processing, the Zymo dataset contained 69,367 sequences with a length ranging from 1,187 to 1,518 bps.

### 3.2 Experimental Setting

We compared our method with seven other alignment-free methods, namely, *k*-mer [13], ACS [16], Kr [39], FFP [14], kmacs [18], SENSE [20], and NeuroSEED [21]. When evaluating an alignment-free method, estimation accuracy and computational efficiency are the two major considerations. Thus, we calculated the mean relative error (MRE) between alignment distances and distances estimated by an alignment-free method and recorded its computational time. By definition, MRE can be interpreted as the percentage of estimated distances that deviate from alignment distances. Since it is computationally infeasible to align all sequence pairs in a dataset, we randomly sampled 1,000 sequences without replacement and calculated the alignment distances of all 499,500 possible sequence pairs. The Needleman-Wunsch algorithm [31] with the default setting was used for sequence alignment (match score = 2, mismatch score = −3, gap opening score = −5, gap extension score = −2). To minimize random variations, the above sampling process was repeated ten times. Therefore, we generated ten test datasets from each sequence dataset to evaluate the eight competing methods.

**Table 2:**
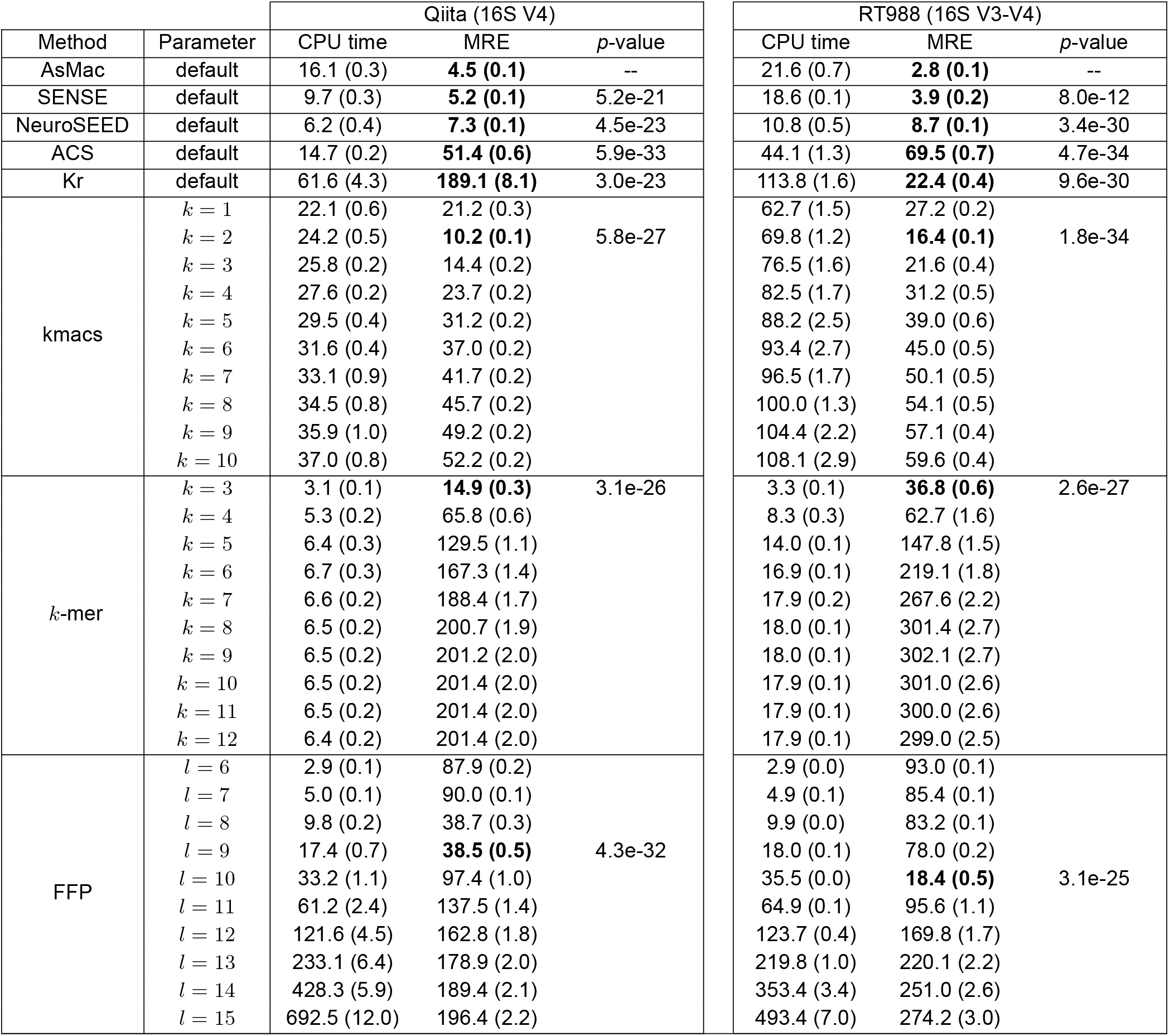
CPU time (seconds) and MRE (%) of eight methods tested on Qiita and RT988 datasets. The numbers in parentheses are standard deviations. When different parameters were used for a given method, the best result is boldfaced. A *p*-value was computed by comparing the MRE result of AsMac with the best result of a competing method.

For kmacs, FFP (V3.19), Kr (V2.0.2), and NeuroSEED, we used the source code provided by the associated articles. Since kmacs is an extension of ACS, we used the kmacs binary by setting *k* = 0 as the implementation of the ACS method. For *k*-mer, we used our C++ implementation, which is optimized for amplicon sequence data by using a sparse *k*-mer count representation. For *k*-mer, kmacs and FFP, different parameters can be applied. Since there is no principal way to estimate the optimal parameter, we tested ten parameters for each method and recorded the best result. For SENSE, we used the default parameters. For NeuroSEED, we used the settings reported to achieve the best results in the original article. For AsMac, we set the number of pattern filters to 300 and the pattern length to 20. For AsMac, SENSE, and NeuroSEED, training of a model for each sequence dataset is required. Since SENSE can only be applied to sequences of roughly equal length, it was trained and tested only on the Qiita and RT988 datasets. To form a training dataset, we randomly sampled sequences from a dataset without replacement and calculated the alignment distances of 5 million sequence pairs. To prevent information leakage, we verified that none of the sequences used for training were included in the test data. The Adam optimizer [40] was employed to train the Siamese neural networks in SENSE and AsMac, where the learning rate was set to 1e-4 and the number of training epochs was set to 200. All experiments were performed on a 3.3 GHz Quad-Core Intel Core i5 with 16GB memory.

**Figure 2:**
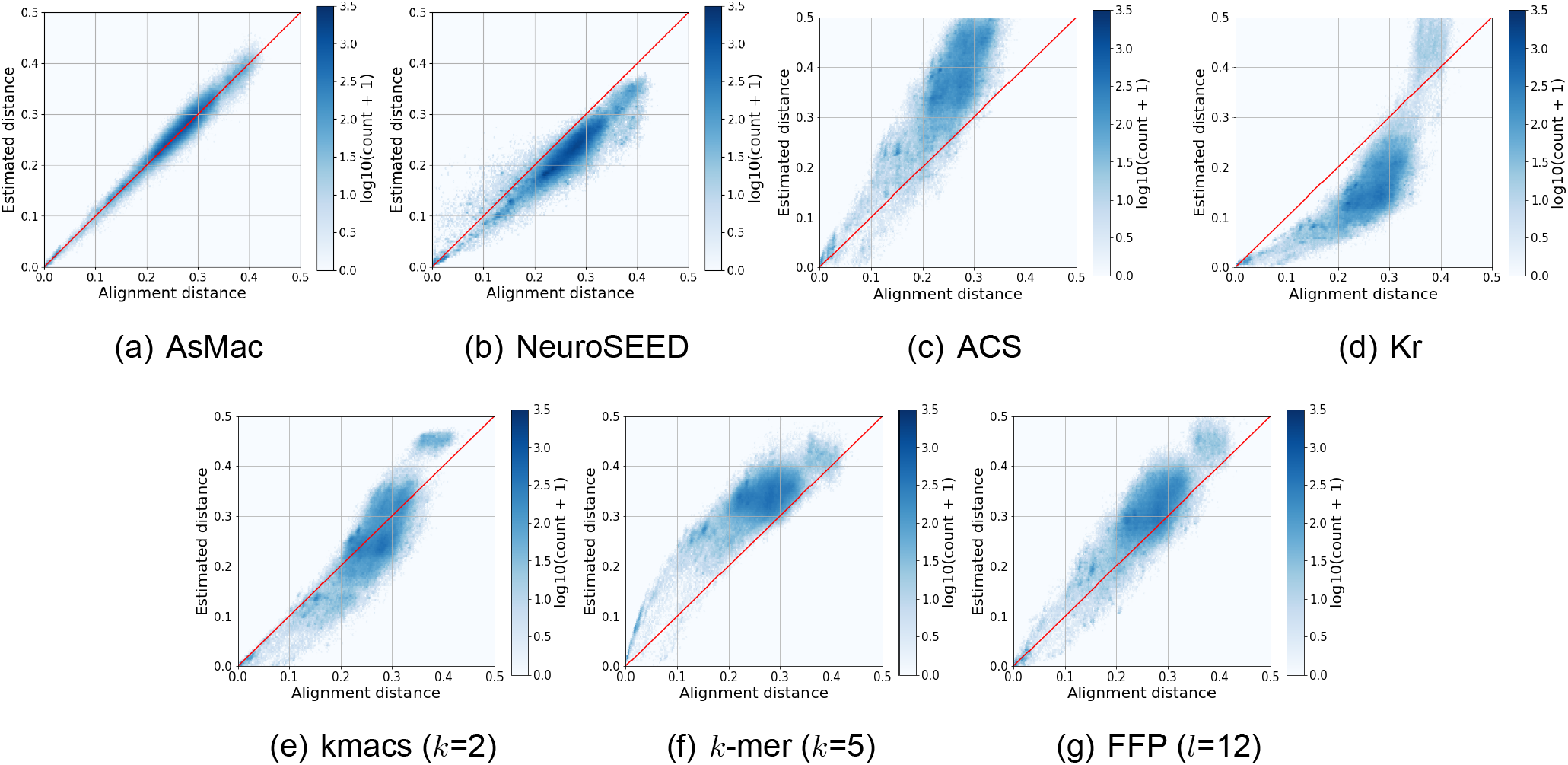
Visualization of alignment distances versus estimated distances computed by seven methods performed on the Greengenes dataset. Each dot represents a sequence pair, and the color of a hex bin represents the number of sequence pairs in the bin. The number in a parenthesis is the parameter that achieved the best result for the corresponding method.

### 3.3 Benchmark Study of Accuracy and Efficiency

First, we applied the eight methods to the Qiita and RT988 datasets, in which sequences cover only a sub-region of the 16S rRNA gene and have similar lengths. Table 2 reports the MREs and CPU time of the eight methods, obtained by averaging over ten runs, and Supplementary Figs. 1 and 2 present the estimated distances against the alignment distances for the two datasets, respectively. In terms of prediction accuracy, AsMac performed slightly better than SENSE and NeuroSEED and significantly outperformed the best results of all other methods by a large margin on both datasets. In terms of running time, while *k*-mer is the fastest algorithm, its performance is not comparable to our method.

**Figure 3:**
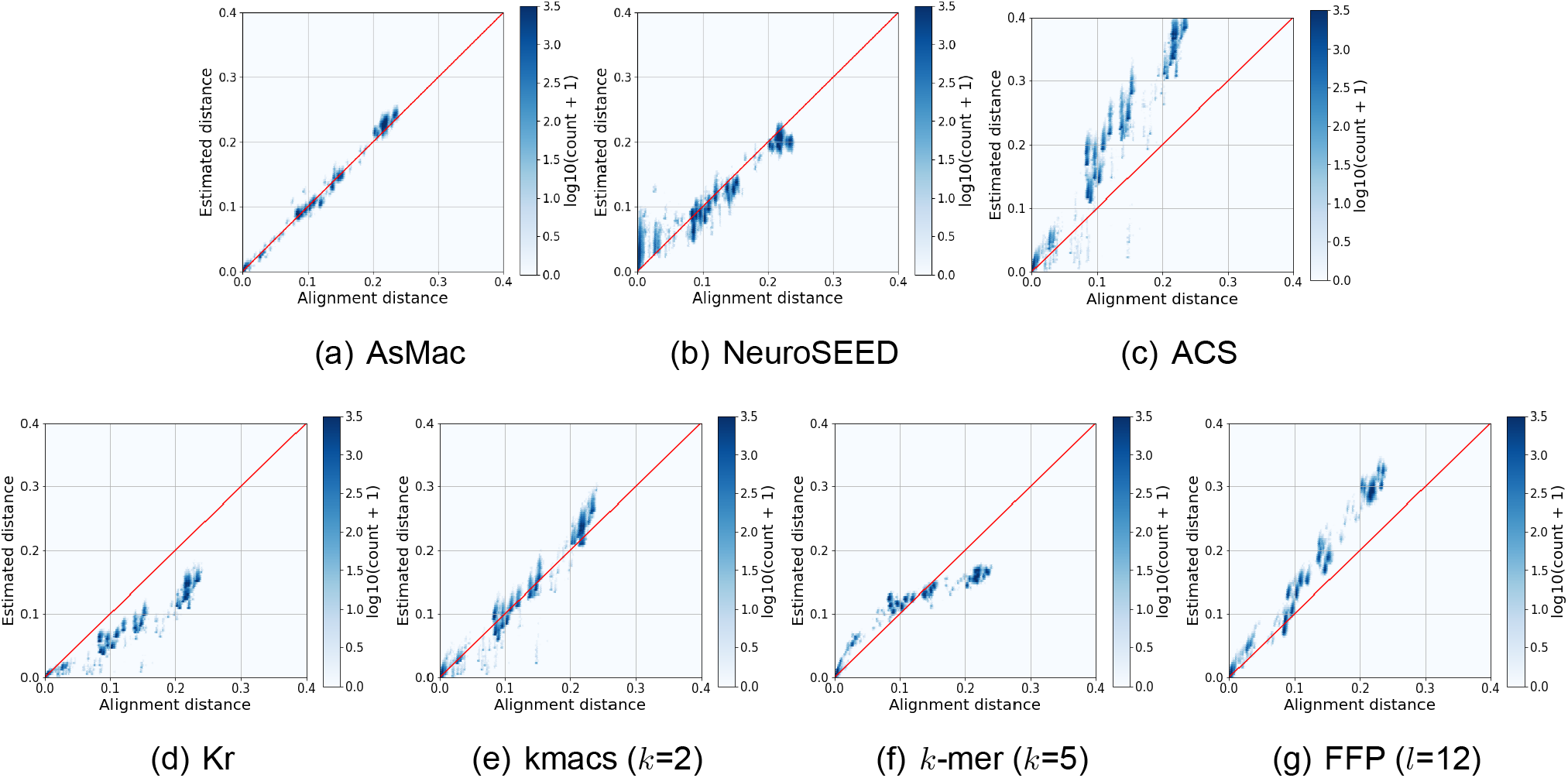
Visualization of alignment distances versus estimated distances computed by seven methods performed on the Zymo dataset. For AsMac and NeuroSEED, a model was trained on the Greengenes dataset and applied to the Zymo dataset.

Next, we applied the competing methods to the Greengenes, Silva-23S, and Labonte lake datasets, where the sequences are of variable lengths. Since there is no biologically meaningful way to trim sequences to an equal length, SENSE cannot be used. Table 3 reports the MREs and CPU time of the seven tested methods. Fig. 2, Supplementary Fig. 3 and 4 present the estimated distances against the alignment distances for the three datasets, respectively. Similar to the results obtained on the Qiita and RT988 datasets, the prediction accuracies of AsMac were significantly better than the best results of all other competing methods.

Finally, we tested the generalization capability of AsMac and NeuroSEED by training a model using the Greengenes dataset and applying it to the Zymo dataset. The results are reported in Table 3 and Fig. 3. Again, AsMac significantly outperformed the other approaches. We noted that, compared to the result obtained from the Greengenes dataset, the prediction accuracy of NeuroSEED declined dramatically. This is likely to be because the Zymo dataset contains a large number of similar sequences that differ by only a few indels, and the CNN structure used by NeuroSEED does not accommodate indels well. As shown in Fig. 3(b), NeuroSEED severely overestimated the pairwise distances of similar sequences. In contrast, by using the ASM structure, our method maintained a high level of accuracy in predicting alignment distances in this situation.

**Table 3:**
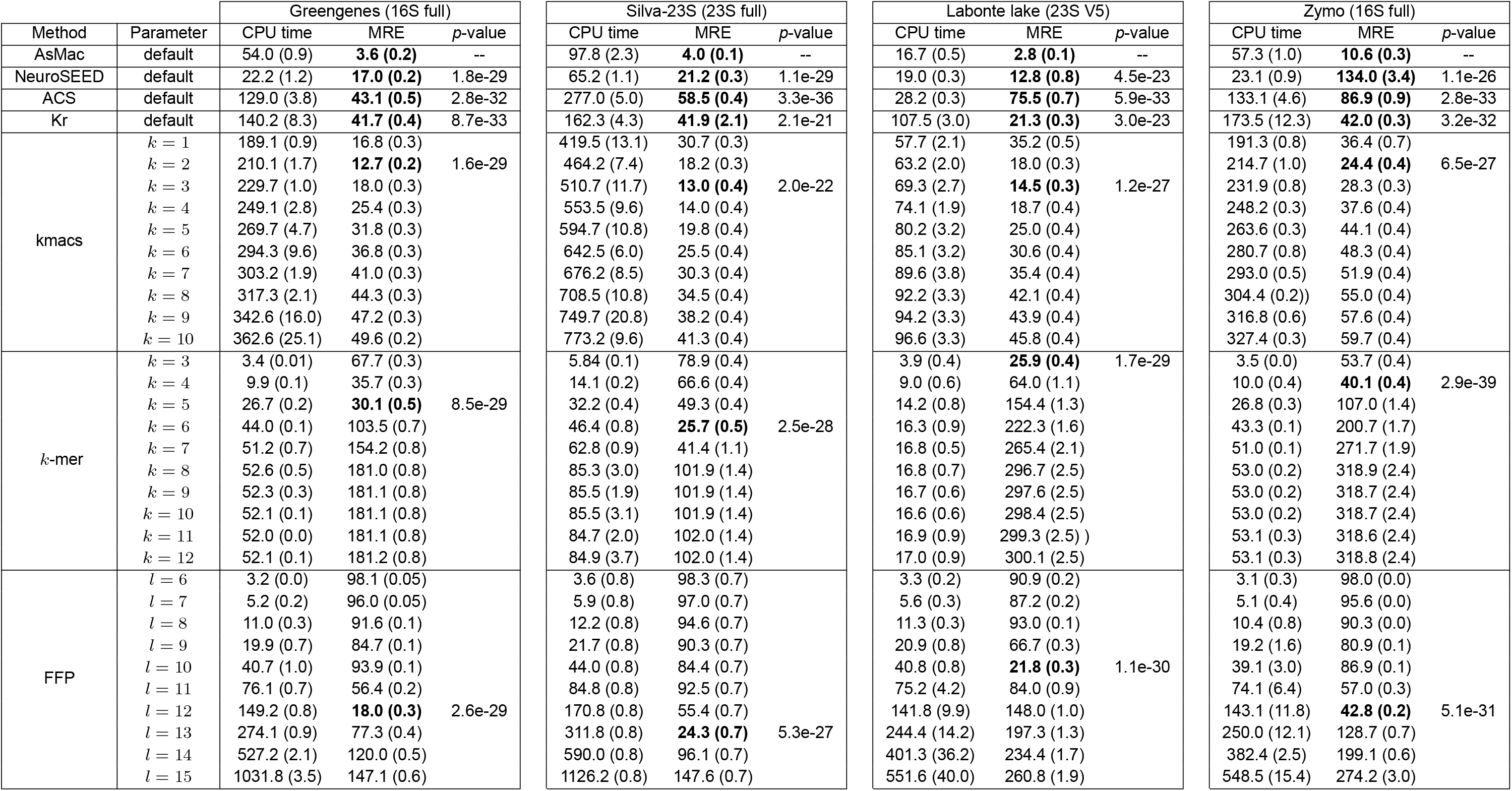
CPU time (seconds) and MRE (%) of seven methods tested on Greengenes, Silva-23S, Labonte lake, and Zymo datasets.

**Figure 4:**
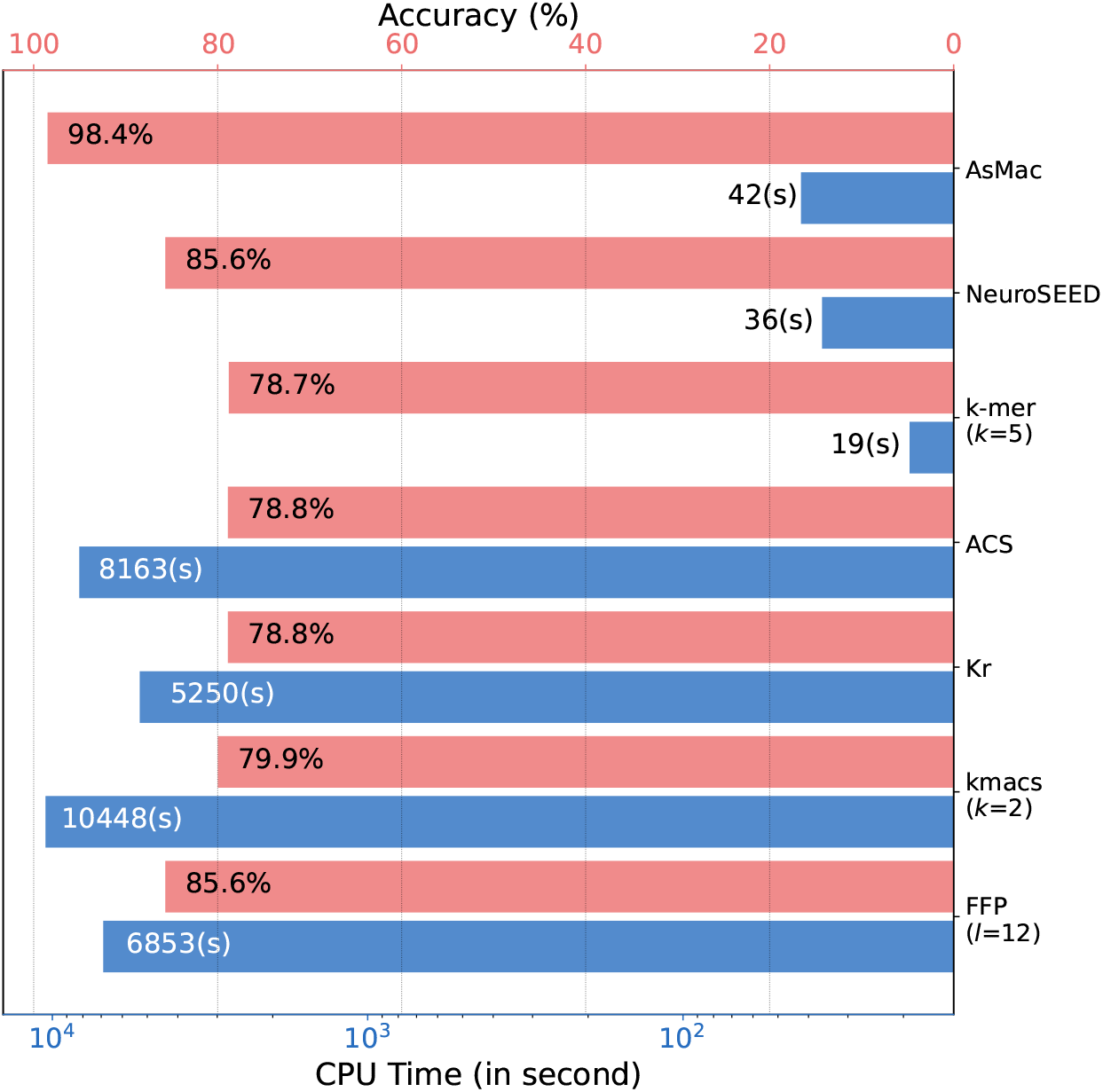
Comparison of taxonomy prediction accuracy and CPU time of seven methods.

### 3.4 Taxonomy Assignment

To further evaluate the effectiveness and utility of the proposed method, we conducted an experiment where we used our approach to perform taxonomy assignment of a given sequence dataset by searching against a reference database. For the purpose of this study, we constructed a query dataset by randomly sampling 3,000 sequences from the Zymo dataset. We used as the reference database the GG97 dataset, a subset of the Greengenes database [35]. This is the default closed-reference database used by QIIME [4] and contains 30,592 sequences with complete taxonomy annotations at the genus level. We performed a database search using the NW algorithm and annotated each query sequence using the genus of the best-matched reference sequence. We used the result obtained by the NW algorithm as the ground truth and compared the performance of our method with the six competing methods used in the Greengenes study. For kmacs, *k*-mer and FFP, the parameters were set to be those that achieved the best performance in the Greengenes study. All methods were performed in parallel using 4 threads.

Fig. 4 reports the annotation accuracy and CPU time of the seven methods tested. Our method achieved 98.4% prediction accuracy, outperforming FFP and NeuroSEED by ~13%, and *k*-mer, ACS, Kr, and kmacs by ~20%. We noticed that kmac did not perform as well as that in the Greengenes and Zymo studies (see Table 3). This may be because for sequence alignment we are concerned about the accuracy of distance estimation for *all* sequence pairs, whereas for sequence annotation we are only interested in sequence pairs that are similar. In terms of computational efficiency, AsMac performed comparably with *k*-mer and NeuroSEED, but ran two orders of magnitude faster than Kr, ACS, kmacs, and FFP. It is worth noting that, compared to the results reported in Table 2, AsMac, NeuroSEED, and *k*-mer ran much faster than the other methods. This is because, for the above three methods, the embedding vectors of the reference sequences can be pre-computed, while for Kr, ACS, and kmacs, the pairwise sequence comparison can only be performed in the presence of both query and reference sequences. Although FFP can pre-process the reference sequences, it is computationally expensive to compute the Kullback-Leibler distances between the query and reference sequences.

**Figure 5:**
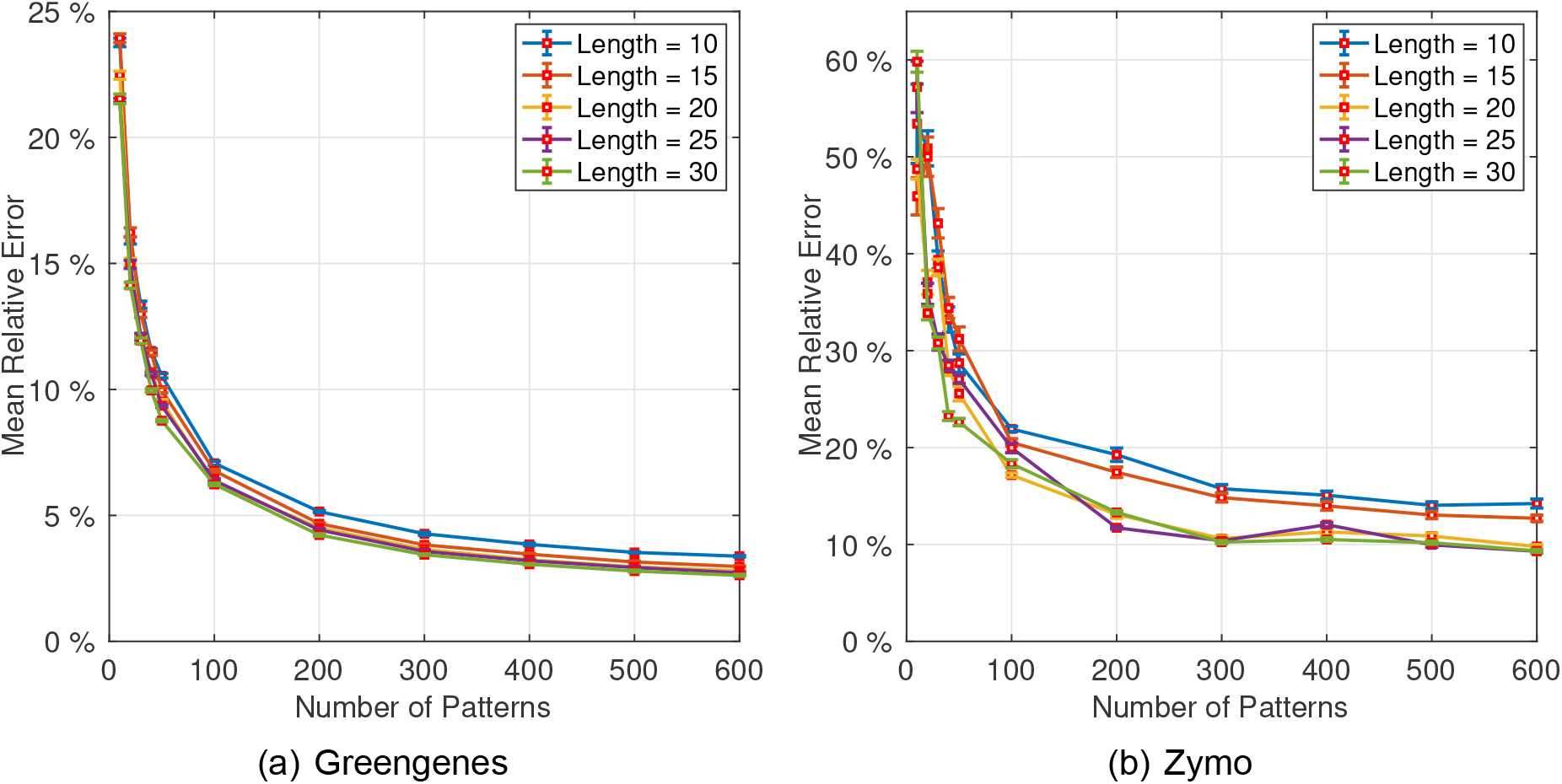
Parameter sensitivity analysis of the proposed method performed on (a) Greengenes and (b) Zymo datasets.

### 3.5 Parameter Sensitivity Analysis

The proposed method has two parameters, namely, the pattern number and the pattern length. We performed a parameter sensitivity analysis to investigate how the method performs with respect to the two parameters. We applied the method to the Greengenes and Zymo datasets and reported in Fig. 5 the MRE results obtained by using various pattern numbers and lengths. We can see that, with an increase of the number of patterns, the prediction errors dropped quickly and then flattened when the pattern number was larger than 300. We also observed that our method achieved similar results once the pattern length was larger than 20. For a balance between accuracy and efficiency, we set the pattern number to 300 and the pattern length to 20 as the default parameters.

## 4 Discussion and Conclusion

In this paper, we described a new neural network-based method for alignment-free sequence comparison. To demonstrate the effectiveness of the proposed method, we conducted a large-scale benchmark study using prokaryotic rRNA sequence datasets. The study showed that the new method performs much more accurately than its data-independent counterparts, and that the method is robust against the presence of deletions and insertions and can handle sequences of varying lengths. The taxonomy assignment experiment further demonstrated the potential utility of our method for practical applications. Compared to data-independent methods, a major drawback of our approach is that it requires training. We should emphasize that *all* supervised learning-based approaches suffer from this drawback. For example, the speech-recognition system used in smartphones requires training, which may take weeks. However, the system installed in smartphones is a trained model, which can perform speech recognition in a matter of milliseconds. To address the aforementioned drawback, we have published five trained neural-network models for some commonly used prokaryotic marker genes (16S rRNA gene, 23S rRNA gene, 16S V3 region, 16S V3-V4 region, and 23S V5 region) in an accompanying website that researchers can use directly to process their own datasets. We should point out that the models for 16S rRNA and 23S rRNA genes were trained on the sequences obtained from databases and may have a better generalization capability than the other models that were built by using the sequences obtained from specific studies. In future work, we will continue to refine the models by using sequence data from diverse studies. We also plan to build models for other marker sequences (e.g., the 18S rRNA gene and internal transcribed spacer sequences of eukaryotes). We will explore the use of ASM for local sequence comparisons, and the aggregation of the ASM structure to form a multi-layer structure, which may further improve prediction accuracy. It will also be of interest to perform in-depth analyses to investigate what type of information in nucleotide sequences has been encoded by the patterns in a neural-network model.

## Acknowledgments

The authors thank the Associate Editor Dr. Aida Ouangraoua and the anonymous reviewers for their valuable suggestions. This work is supported in part by funds from the National Institutes of Health [R01CA241123 to Y. S. and S. G., R01CA266113 to Y. S. and S. G.].

## References

[1] John C Wooley, Adam Godzik, et al. A primer on metagenomics. PLoS Computational Biology, 6(2):e1000667, 2010.

[2] Jack A Gilbert, Robert A Quinn, et al. Microbiome-wide association studies link dynamic microbial consortia to disease. Nature, 535(7610):94--103, 2016.

[3] Editorial. Your microbes, your health. Science, (342):1440--1441, 2013.

[4] J Gregory Caporaso, Justin Kuczynski, et al. QIIME allows analysis of high-throughput community sequencing data. Nature Methods, 7(5):335--336, 2010.

[5] Robert C Edgar. Search and clustering orders of magnitude faster than BLAST. Bioinformatics, 26(19):2460--2461, 2010.

[6] Yunpeng Cai and Yijun Sun. ESPRIT-Tree: hierarchical clustering analysis of millions of 16S rRNA pyrosequences in quasilinear computational time. Nucleic Acids Research, 39(14):e95, 2011.

[7] Yunpeng Cai, Wei Zheng, et al. ESPRIT-Forest: parallel clustering of massive amplicon sequence data in subquadratic time. PLOS Computational Biology, 13(4):e1005518, 2017.

[8] Wei Zheng, Qi Mao, et al. A parallel computational framework for ultra-large-scale sequence clustering analysis. Bioinformatics, 35(3):380--388, 2019.

[9] Andrzej Zielezinski, Susana Vinga, et al. Alignment-free sequence comparison: benefits, applications, and tools. Genome Biology, 18(1):186, 2017.

[10] Oliver Bonham-Carter, Joe Steele, et al. Alignment-free genetic sequence comparisons: a review of recent approaches by word analysis. Briefings in Bioinformatics, 15(6):890--905, 2014.

[11] Kai Song, Jie Ren, et al. New developments of alignment-free sequence comparison: measures, statistics and next-generation sequencing. Briefings in Bioinformatics, 15(3):343--353, 2014.

[12] Andrzej Zielezinski, Hani Z Girgis, et al. Benchmarking of alignment-free sequence comparison methods. Genome Biology, 20:144, 2019.

[13] Samuel Kariin and Chris Burge. Dinucleotide relative abundance extremes: a genomic signature. Trends in Genetics, 11(7):283--290, 1995.

[14] Gregory E Sims, Se-Ran Jun, et al. Alignment-free genome comparison with feature frequency profiles (FFP) and optimal resolutions. Proceedings of the National Academy of Sciences, 106(8):2677--2682, 2009.

[15] Lei Gao and Ji Qi. Whole genome molecular phylogeny of large dsDNA viruses using composition vector method. BMC Evolutionary Biology, 7(1):1--7, 2007.

[16] Igor Ulitsky, David Burstein, et al. The average common substring approach to phylogenomic reconstruction. Journal of Computational Biology, 13(2):336--350, 2006.

[17] Bernhard Haubold, Peter Pfaffelhuber, et al. Estimating mutation distances from unaligned genomes. Journal of Computational Biology, 16(10):1487--1500, 2009.

[18] Chris-Andre Leimeister and Burkhard Morgenstern. Kmacs: the k-mismatch average common substring approach to alignment-free sequence comparison. Bioinformatics, 30(14):2000--2008, 2014.

[19] Yang Gao and Liaofu Luo. Genome-based phylogeny of dsDNA viruses by a novel alignment-free method. Gene, 492(1):309--314, 2012.

[20] Wei Zheng, Le Yang, et al. SENSE: Siamese neural network for sequence embedding and alignment-free comparison. Bioinformatics, 35(11):1820--1828, 2019.

[21] Gabriele Corso, Zhitao Ying, et al. Neural distance embeddings for biological sequences. In Advances in Neural Information Processing Systems, pages 1--12, 2021.

[22] Yann LeCun, Yoshua Bengio, et al. Deep learning. Nature, 521(7553):436--444, 2015.

[23] Martin Sundermeyer, Ralf Schlüter, et al. LSTM neural networks for language modeling. In INTERSPEECH, 2010.

[24] Qiang Wang, Bei Li, et al. Learning deep transformer models for machine translation. arXiv preprint arXiv:1906.01787, 2019.

[25] Jane Bromley, Isabelle Guyon, et al. Signature verification using a “siamese” time delay neural network. In Advances in Neural Information Processing Systems, pages 737--744, 1993.

[26] Peter H Sellers. The theory and computation of evolutionary distances: pattern recognition. Journal of Algorithms, 1(4):359--373, 1980.

[27] Satoshi Koide, Keisuke Kawano, et al. Neural edit operations for biological sequences. In Advances in Neural Information Processing Systems, pages 4960--4970, 2018.

[28] Marco Cuturi and Mathieu Blondel. Soft-DTW: a differentiable loss function for time-series. In International Conference on Machine Learning, pages 894--903, 2017.

[29] Vinod Nair and Geoffrey E Hinton. Rectified linear units improve restricted Boltzmann machines. In International Conference on Machine Learning, pages 807--814, 2010.

[30] Adam Paszke, Sam Gross, et al. Pytorch: An imperative style, high-performance deep learning library. In Advances in Neural Information Processing Systems, pages 8026--8037, 2019.

[31] Saul B Needleman and Christian D Wunsch. A general method applicable to the search for similarities in the amino acid sequence of two proteins. Journal of Molecular Biology, 48(3):443--453, 1970.

[32] Jose C Clemente, Erica C Pehrsson, et al. The microbiome of uncontacted Amerindians. Science Advances, 1(3):e1500183, 2015.

[33] Robert J Genco, Michael J LaMonte, et al. The subgingival microbiome relationship to periodontal disease in older women. Journal of Dental Research, 98(9):975--984, 2019.

[34] Blaire Steven, Sage McCann, et al. Pyrosequencing of plastid 23s rrna genes reveals diverse and dynamic cyanobacterial and algal populations in two eutrophic lakes. FEMS Microbiology Ecology, 82(3):607--615, 2012.

[35] Daniel McDonald, Morgan N Price, et al. An improved Greengenes taxonomy with explicit ranks for ecological and evolutionary analyses of bacteria and archaea. The ISME Journal, 6(3):610, 2012.

[36] Elmar Pruesse, Christian Quast, et al. Silva: a comprehensive online resource for quality checked and aligned ribosomal RNA sequence data compatible with ARB. Nucleic Acids Research, 35(21):7188--7196, 2007.

[37] Benjamin J Callahan, Joan Wong, et al. High-throughput amplicon sequencing of the full-length 16S rRNA gene with single-nucleotide resolution. Nucleic Acids Research, 47(18):e103, 2019.

[38] Benjamin J Callahan, Paul J McMurdie, et al. DADA2: high-resolution sample inference from Illumina amplicon data. Nature Methods, 13(7):581--583, 2016.

[39] Mirjana Domazet-Lošo and Bernhard Haubold. Efficient estimation of pairwise distances between genomes. Bioinformatics, 25(24):3221--3227, 2009.

[40] Diederik P Kingma and Jimmy Ba. Adam: A method for stochastic optimization. In Inter-national Conference on Learning Representations, pages 1--13, 2014.

