## Supplementary figures for "Alignment-free Comparison of Metagenomics Sequences via Approximate String Matching"

### Supplementary Data

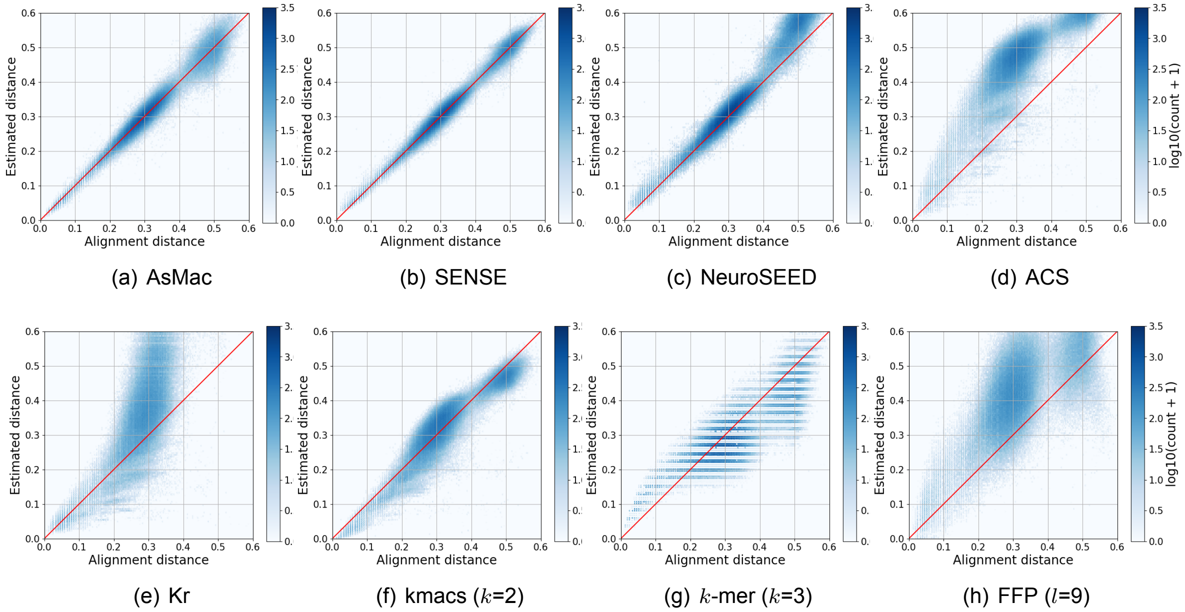


(a) AsMac (b) SENSE (c) NeuroSEED (d) ACS

(e) Kr (f) kmacs ($k$=2) (g) $k$-mer ($k$=3) (h) FFP ($l$=9)

Figure 1: Visualization of alignment distances versus estimated distances computed by eight methods performed on the Qiita dataset. Each dot represents a sequence pair, and the color of a hex bin represents the number of sequence pairs in the bin. The number in a parenthesis is the parameter that achieved the best result for the corresponding method.

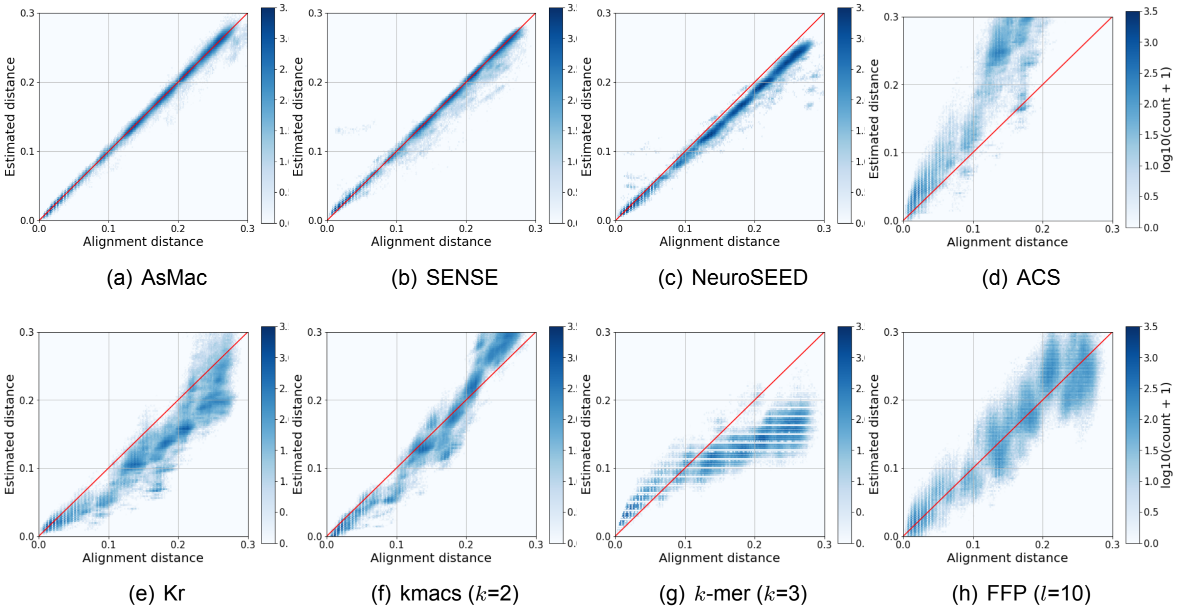


(a) AsMac (b) SENSE (c) NeuroSEED (d) ACS

(e) Kr (f) kmacs ($k$=2) (g) $k$-mer ($k$=3) (h) FFP ($l$=10)

Figure 2: Visualization of alignment distances versus estimated distances computed by eight methods performed on the RT988 dataset.

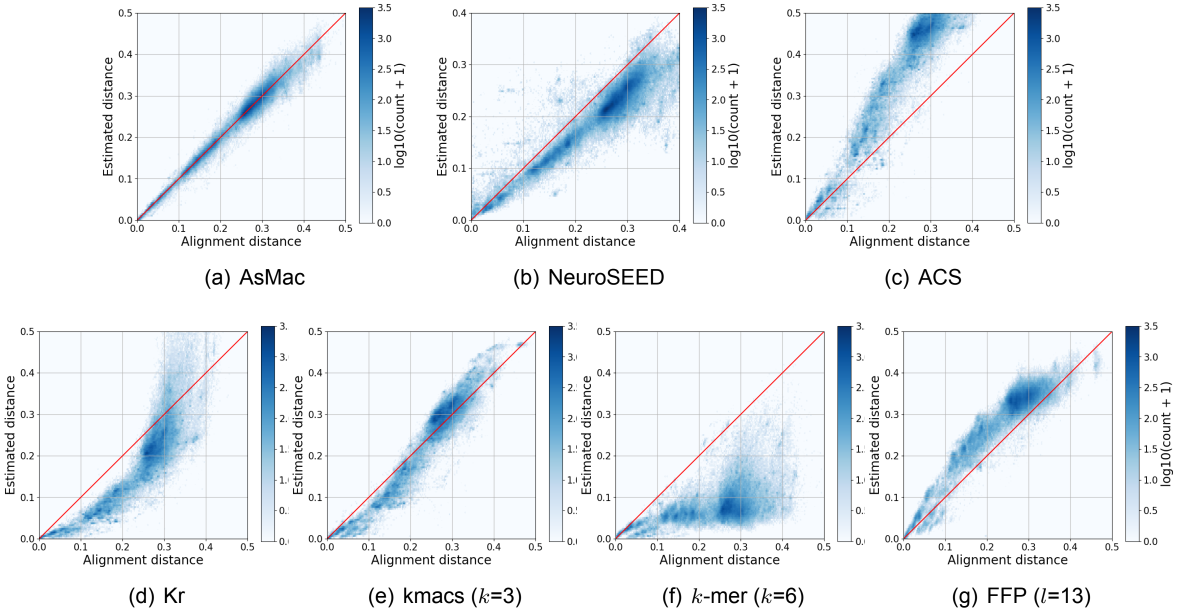


(a) AsMac (b) NeuroSEED (c) ACS

(d) Kr (e) kmacs ($k$=3) (f) $k$-mer ($k$=6) (g) FFP ($l$=13)

Figure 3: Visualization of alignment distances versus estimated distances computed by seven methods performed on the Silva-23S dataset.

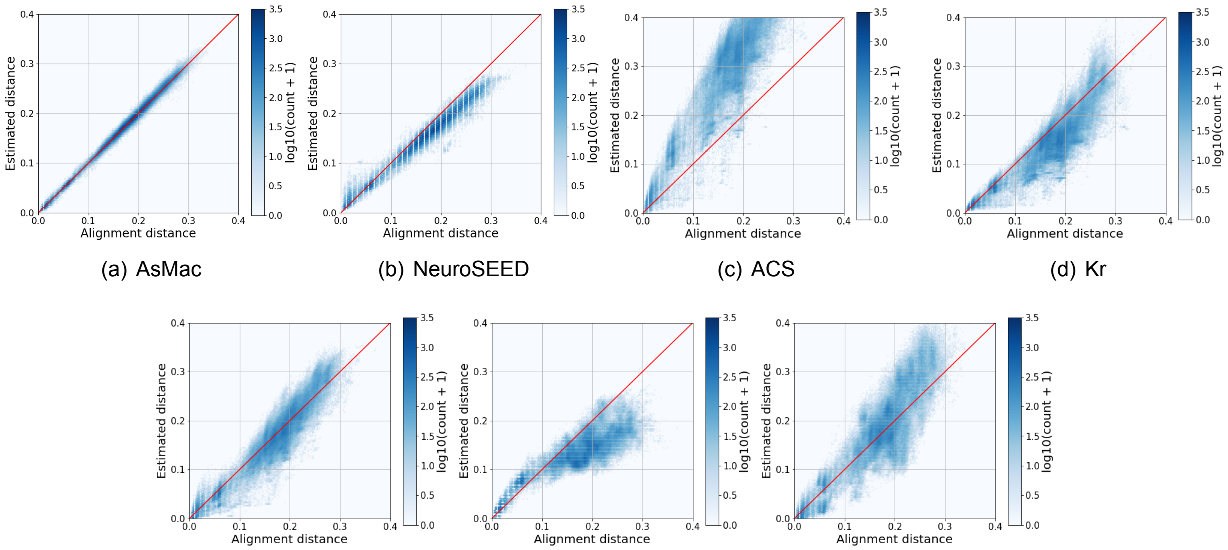


(a) AsMac (b) NeuroSEED (c) ACS (d) Kr

(e) kmacs ($k$=2) (f) $k$-mer ($k$=5) (g) FFP ($l$=12)

Figure 4: Visualization of alignment distances versus estimated distances computed by seven methods performed on the Labonte lake dataset.
